# A model for the regulation of the timing of cell division by the circadian clock in the cyanobacterium *Synechococcus elongatus*

**DOI:** 10.1101/765669

**Authors:** Po-Yi Ho, Bruno M.C. Martins, Ariel Amir

## Abstract

Cells of the cyanobacterium *Synechococcus elongatus* possess a circadian clock in the form of three core clock proteins (the Kai proteins) whose concentrations and phosphorylation states oscillate with daily periodicity under constant conditions [1]. The circadian clock regulates the cell cycle such that the timing of cell divisions is biased towards certain times during the circadian period [2, 3, 4, 5], but the mechanism underlying how the clock regulates division timing remains unclear. Here, we propose a mechanism in which a protein limiting for division accumulates at a rate proportional to cell volume growth and modulated by the clock. This “modulated rates” model, in which the clock signal is integrated over time to affect division timing, differs fundamentally from the previously proposed “gating” concept, in which the clock is assumed to suppress divisions during a specific time window [2, 3]. We found that while both models can capture the single-cell statistics of division timing in *S. elongatus*, only the modulated rates model robustly places divisions away from darkness during changes in the environment. Moreover, within the framework of the modulated rates model, existing experiments on *S. elongatus* are consistent with the simple mechanism that division timing is regulated by the accumulation of a division limiting protein in phase with genes whose activity peak at dusk.

## 2 Results

### Modeling the growth and division of *S. elongatus* cells

To construct a model to describe how the clock affects division timing, we analyzed data from Ref. [5], which observed the growth and division of single cells for a wild type strain of *S. elongatus* and a strain without the Kai B and Kai C proteins, referred to here as the clock-deletion strain. Since *S. elongatus* cells require light to grow, they were grown and imaged under constant light (LL) or periodic cycles of light and darkness (12:12 LD, i.e. 12 hours of light with a graded intensity profile, followed by 12 hours of darkness, and 16:8 LD) to probe the effects of the clock on division timing under different environments. For each cell, its length at birth *l_b_* and division *l_d_* and its generation time (the time between birth and division) *t_d_* were measured. Before imaging, the cells were grown under 12:12 LD to entrain and synchronize the activity of the clock to the environmental light conditions. The circadian phase *θ* corresponding to the internal, subjective time of day encoded by the clock can then be assumed to be set to the environmental light-dark cycle. We defined *θ* = 0 h to be dawn, or the beginning of the period under light. Each cell can then be assigned a circadian phase at birth *θ_b_*. We analyzed the distributions of (denoted *p* (·)) and correlations among the four stochastic variables (*l_b_*, *l_d_*, *t_d_*, *θ_b_*), and compared these statistics of division timing to those generated by our models (Fig. 1). Similar approaches have led to insights on other aspects of microbial and also eukaryotic cell cycles, including how DNA replication might be coupled to division timing [6, 7, 8, 9, 10, 11, 12, 13, 14, 15].

**Figure 1:**
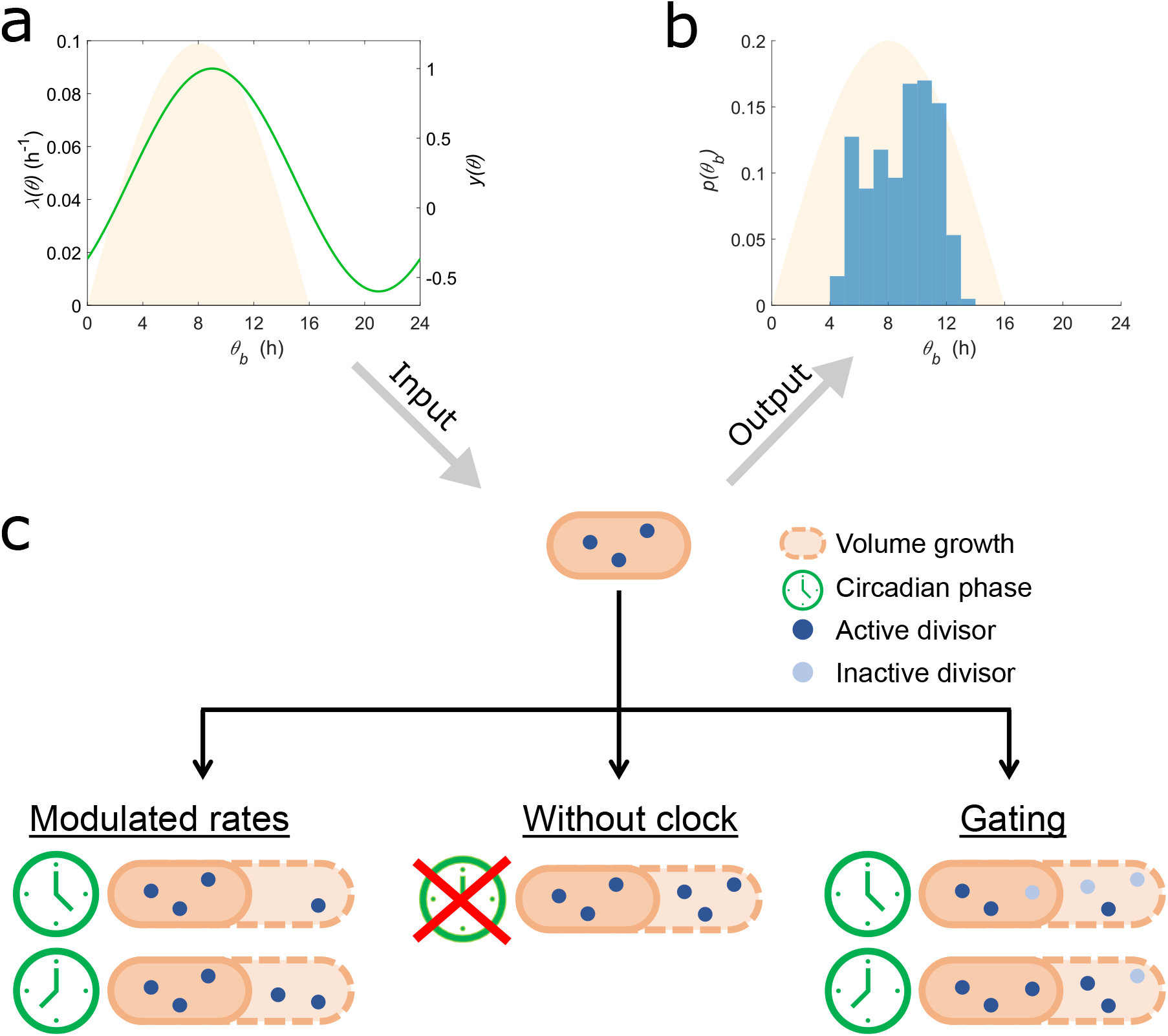
Two models for the regulation of division timing by the circadian clock. Both models take as inputs (a) the environmental light-dark cycles (*λ* (*θ*), yellow shade) and a modulation function (*y* (*θ*), green line) that determines how the clock affects division timing to give as outputs (b) the single-cell distributions of and correlations among cell length at birth *l_b_* and division *l_d_*, the circadian phase at birth *θ_b_*, and the generation time *t_d_*. Shown is an experimentally observed distribution of *θ_b_* for *S. elongtaus* under periodic conditions, showing that divisions occur away from dawn and dusk [5]. (c) (Middle) The divisor accumulation model without the clock, Eq. 3. The divisor is accumulated at a rate proportional to volume growth. (Left) The modulated rates model, Eq. 7. The divisor accumulation rate is modulated by the clock. (Right) The gating model, Eq. 9. The divisor accumulation rate is not affected by the clock, but only a fraction of divisors, determined by the current circadian phase, is active towards reaching the threshold.

We first modeled the growth mode of single cells, which has significant implications for cell cycle regulation (see Discussion) [7, 14]. The growth mode of *S. elongatus* cells can be approximated to be exponential, with a rate dependent on the environmental light intensity [4, 5]. We therefore modeled the growth of volume *V* as

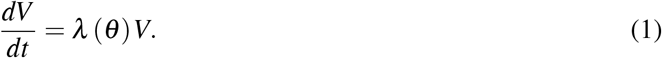

The growth rate *λ* (*θ*) may depend on the light intensity, which is a function of *θ* for the periodic environments under consideration. Under LL, *λ* (*θ*) can be approximated as constant for our purposes, although in reality it varies up to ≈ 5% with the circadian phase [5]. Under LD, *λ* (*θ*) is approximately proportional to the environmental light intensity. The experimental light intensity profile is sinusoidal during the period under light. We therefore model *λ* (*θ*) as

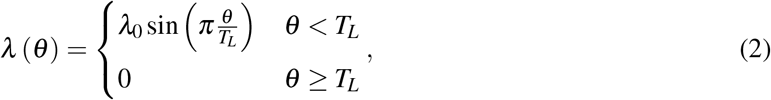

where *λ*_0_ is the maximum growth rate and *T_L_* is the duration of the period under light. *λ*_0_ and *T_L_* are known parameters. For our analyses, we use cell volume and cell length interchangeably since volume can be well approximated as proportional to cell length in rod-shaped bacteria that grow by elongation such as *S. elongatus* [16]. Experimentally, cells divide approximately symmetrically with small fluctuations in the division ratio (i.e. 0.51 ± 0.02 in the data set for wild type cells under LL). We assumed perfectly symmetrical divisions in our models.

### Divisor accumulation can describe division timing in a clock-deletion strain

To construct a basic model of division timing without a clock, we considered the experiments on the clock-deletion strain. Under LL, the clock-deletion strain behaves, with minor deviations, like several other microbes whose cells appear to add a constant size from birth to division on average (Fig. 2a) [6, 8, 11, 12, 17, 18]. Inspired by models that sought to describe such single-cell correlations (SM Section 5.1) [9, 19, 20], we considered the following “divisor accumulation” model. Its basic component is the accumulation of a divisor protein limiting for division, whose amount is denoted by *X*, at a rate proportional to volume growth,

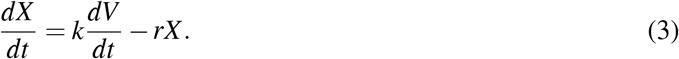

**Figure 2:**
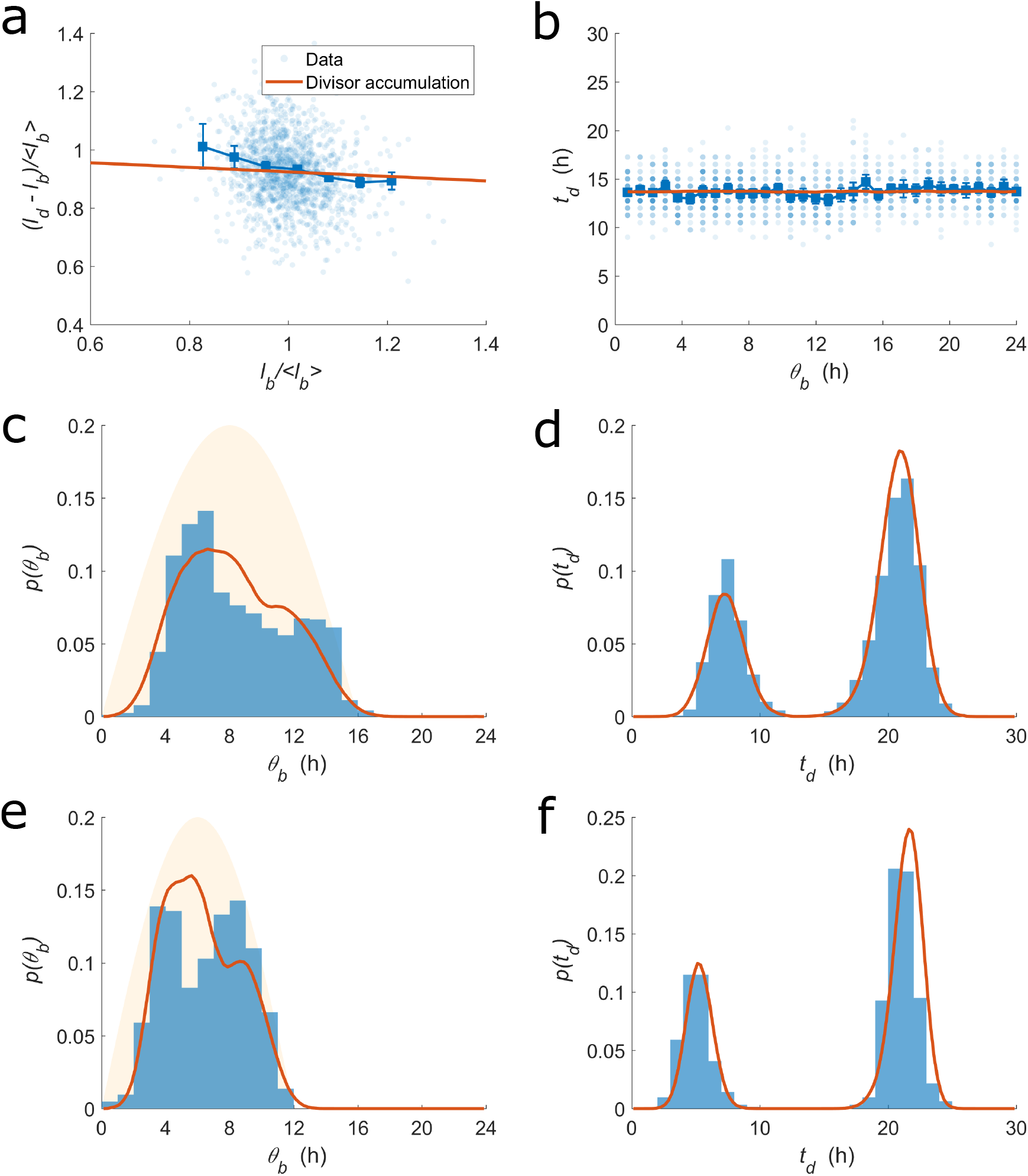
Divisor accumulation can describe division timing in the clock-deletion strain under LL (a,b), and under 16:8 (c,d) or 12:12 (e,f) LD. The correlations (a,b) and distributions (c-f) of the stochastic variables as defined in the caption of Fig. 1. ⟨·⟩ denotes the average over all single-cells. Blue denotes data from Ref. [5]. Red lines denote predictions of the divisor accumulation model. (a,b) Small points represent single-cell data. Large squares are averages binned according to the x-axis, with error bars showing the standard error of the mean. (c,e) Yellow shade shows the shape of the light intensity profile. Table S2 contains the parameter values used.

Here, *r* is the degradation rate of the divisor. Division occurs upon the accumulation of a threshold amount *X*_0_ of divisors. Divisors are consumed during division so that the amount of divisors is zero at birth, denoted by *t* = 0. That is,

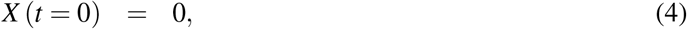

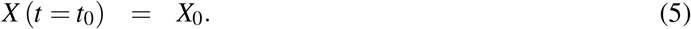

The resetting of divisors could be describing a scenario similar to the disassembly of the divisome in *E. coli* [21]. In Eq. 5, *t*_0_ is the deterministic generation time. On top of the deterministic dynamics of Eqs. 3–5, we implement a time-additive noise to model the stochasticity in division timing due to, for example, noise in gene expression (e.g. Refs. [7, 22]). The stochastic generation time *t_d_* is

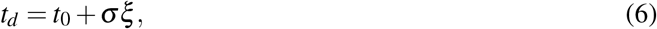

where *ξ* is a normal random variable with zero mean and unit standard deviation, and *σ* is the magnitude of the time-additive noise. In Eqs. 3 and 5, we set *k* = 1 and *X*_0_ = 1 because we were not interested in the absolute magnitudes of the concentration of the divisor or the cell volume. Instead, we analyzed statistics such as coefficient of variations (CV, the standard deviation divided by the mean) and correlations coefficients that are independent of the absolute magnitudes. The free parameters of the model are *r* and *σ*. The best fit value of *r* was determined for the clock-deletion strain under 16:8 LD, which admitted a more precise determination of *r* than other conditions (Methods). The resulting value of *r* was 0.025 ± 0.006 h^−1^, which corresponds to a half life of approximately 28 h, and was assumed to be the same for all other conditions (Methods). *σ* was determined separately for each condition, analogous to the fact that bacterial cells grown under different conditions might exhibit different magnitudes of stochasticity in division timing [8]. Although the model does not specify the molecular identity of the divisor, it might be describing, for example, the accumulation of FtsZ, a protein implicated for cell division in some bacteria [13, 23, 24].

We then compared the divisor accumulation model, Eqs. 1–6, with the experiments on the clock-deletion strain. Under LL, the model predicts close to no correlations between *l_d_* − *l_b_* and *l_b_*, in approximate agreement with experiments (Fig. 2a). Moreover, since the model does not contain a clock, *t_d_* is independent of *θ_b_*, again in agreement with experiments (Fig. 2b). Under LD, experiments showed that the value of *p*(*θ_b_*) is small near dawn. The model captures this observation because the divisors degrade, so that cells typically do not have enough divisors to divide immediately after dawn (Fig. 2ce, SM Section 5.1). Under LD, *p*(*t_d_*) is bimodal because some cells divide before reaching a period of darkness (short-generation cells), whereas other cells must wait through a period of darkness before division (long-generation cells). The model is able to capture the mean generation times of both short- and long-generation cells (Fig. 2df, SM Section 5.1). Moreover, the model predictions for the distributions of and the correlations between the other stochastic variables also agree with experiments (Fig. S3abc). Taken together, divisor accumulation is a simple mechanistic model that can capture the statistics of division timing in the clock-deletion strain.

### Divisor accumulation with modulated rates can describe division timing with a circadian clock

To construct a mechanistic model for how the clock affects division timing, we incorporated the effects of the clock into the divisor accumulation model, and compared the resulting model with the experiments on the wild type strain. Under LL, the clock generates correlations between *θ_b_*, *l_b_*, and *t_d_* that cannot be captured by the divisor accumulation model without a clock (Fig. 3ab). We therefore considered a modulated rates model where the rate of accumulation of the divisor is modulated by a periodic function *y* (*θ*),

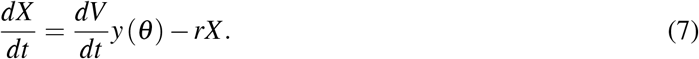

*y* (*θ*) could be describing, for example, the approximately sinusoidal promoter activity of FtsZ under LL [25]. We therefore assumed the following sinusoidal form,

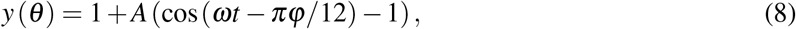

where *ω* = 2*π*/ (24 h), and *A* and *φ* are the magnitude and phase offset of the modulation. The sinusoidal form is reasonable also under periodic LD conditions because the promoter activity of the Kai proteins, and presumably FtsZ, remains sinusoidal during the day [26]. We chose *y* (*θ*) to have a maximum of one, since the absolute magnitude of *y* (*θ*) does not affect the statistics of division timing. We also enforced *X* ≥ 0. We determined the values of the free parameters *A* and *φ* for each condition (Methods, SM Section 5.2), reflecting the fact that the molecular players that implement *y* (*θ*) may depend on environmental light conditions [27], which we discuss in detail below. We also assumed that clocks are entrained quickly relative to the duration of the experiments, so that the measured statistics approximate those for cells entrained under imaging conditions, even though the actual protocol entrained the cells under 12:12 LD even for the 16:8 LD imaging condition.

**Figure 3:**
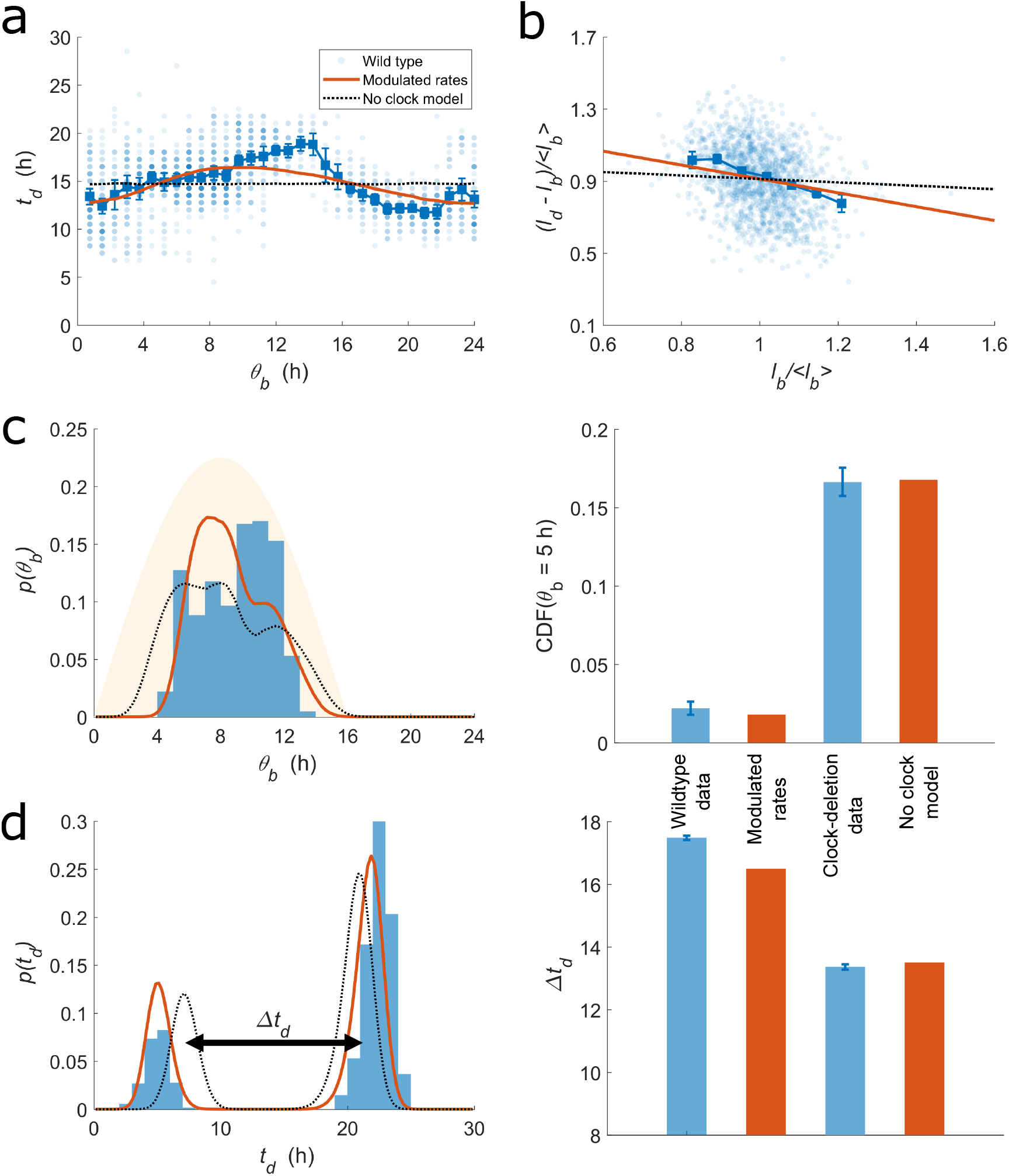
Divisor accumulation with modulated rates can describe division timing in the wild type strain under LL (a,b) and under 16:8 LD (c,d). Figure legends and axes labels are the same as in Fig. 2 except that red lines here denote predictions of the modulated rates model. Dashed black lines denote predictions of modulated rates model with *y* (*θ*) = 1, equivalent to the divisor accumulation model, under the corresponding conditions. (c) The bar plot shows the cumulative fraction of divisions that have occured before the specified circadian phase five hours after dawn. (d) The bar plot shows the difference in the mean generation times of the short- and long-generation cells, Δ*t_d_*. Error bars in (c,d) show the standard deviation of the estimates due to sampling error calculated using bootstrapping. The results under 12:12 LD are shown in Fig. S4. The results for the gating model are shown in Fig. S5. Table S2 contains the parameter values used.

Despite its simplicity, the modulated rates model can capture the correlations between *t_d_* and *θ_b_* under LL (Fig. 3a). The model without further adjustments also captures the correlations between *l_d_* − *l_b_* and *l_b_* (*ρ* = −0.32 ± 0.05, Pearson correlation coefficient with 95% confidence interval; Fig. 3b), which is more negative than in the clock-deletion strain (*ρ* = −0.21 ± 0.05). Such correlations arise because, within the model, cells that are larger at birth likely have just grown through periods where the divisor accumulation rate was repressed by *y* (*θ*), and will therefore tend to grow through periods of derepressed divisor accumulation. The size increments between birth and division of larger cells will therefore be smaller than average (SM Section 5.1). The model also captures the other statistics of division timing (Fig. S3d). In particular, because the statistics were not collected over lineages but over growing populations, *p*(*θ_b_*) is not the same as the distribution of circadian phases at division *p*(*θ_d_*), where *θ_d_* is defined as (*θ_b_* + *t_d_*) mod 24. The model captures both distributions after taking into account the details of the ensemble (SM Section 5.3). Moreover, the model also captures correlations between distantly related cells such as the cousin-cousin correlations between generation times (SM Section 5.4).

The model can also describe the division timing of the wild type strain under LD (Fig. 3cd, Fig. S3ef). Specifically, it captures that wild type cells, compared to cells of the clock-deletion strain, began to divide later after dawn, and stopped dividing sooner before dusk (Fig. 3c). Within the model, divisions are biased to occur away from darkness because *y* (*θ*) peaks near the mid-point of the light period (Fig. 1a). The model also predicts that the clock will decrease (increase) the mean generation time of the short- (long-) generation cells, in agreement with experiments (Fig. 3d). In summary, divisor accumulation with modulated rates, Eq. 7, is a model with two free parameters (*A* and *φ*) that can describe the statistics of division timing in wild type *S. elongatus* under both constant and periodic environments (Fig. 3 and Fig. S3).

### The modulated rates model robustly places divisions away from darkness, whereas the gating model does not

The modulated rates model, in which the clock signal is integrated over time to affect division timing, differs fundamentally from the widely considered gating hypothesis, which assumes that the clock suppresses divisions during a specific time window [2, 3]. We next sought to distinguish between the two hypotheses by incorporating the gating hypothesis into the framework of the divisor accumulation model, and comparing the predictions of the two models. In our gating model, divisors accumulate without modulation by the clock, as in Eq. 3. However, only a fraction *y* (*θ*) of the accumulated divisors is active in contributing to reaching the threshold. That is,

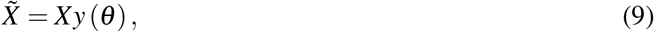

where 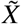 is the amount of active divisors. A threshold amount of active divisors triggers division,

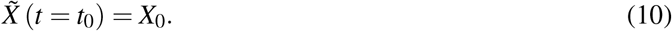

All other aspects of the gating model are the same as the modulated rates model. The gating function *y* (*θ*) could be, for example, a step function equal to zero during the window of suppressed division and one otherwise, which is exactly the case considered in Ref. [3]. To compare the gating and the modulated rates models without additional differences, we considered the case where *y* (*θ*) is sinusoidal as in Eq. 8. By using the same fitting procedure as for the modulated rates model, we found that the gating model can also capture the effects of the clock on the statistics of division timing (Fig. S5). To more clearly distinguish between the two hypotheses, we next sought to understand how the two models differ qualitatively.

First, the best fit values of *φ* suggest that the effect on division timing by the clock is implemented by different molecular players in the two models. For the modulated rates model, one mechanistic interpretation is that *y* (*θ*) describes the promoter activity of the divisor. In this case, the value of *φ* is related to the phase at the peak of the concentration of the divisor (SM Section 5.1). Specifically, we found that the divisor concentration peaks approximately 12 ± 1 hours after dawn under 12:12 LD (Table 1). Experiments have observed that a bioluminescent reporter under the control of the *kaiBC* promoter peaks 14 ± 1 hours after dawn under similar conditions [26]. The approximate agreement between the two suggests that within the modulated rates model, *y* (*θ*) is implemented by molecular players that are roughly in synchrony with expression of the Kai proteins. For the gating model, one mechanistic interpretation is that *y* (*θ*) describes the concentration of an effector that transmits the signal of the clock to affect division timing, since the effector acts immediately to affect the fraction of active divisors. In this case, the best fit values of *φ* in the gating model imply that the effector concentration peaks 8 hours after dawn under 12:12 LD (Table 1). Therefore, within the gating model, *y* (*θ*) would be implemented by molecular players that are not in synchrony with expression of the Kai proteins, in contrast with the modulated rates model. This difference between the two models is reminiscent of the different classes of promoters whose peaking time cluster around either dusk or dawn [28], although the difference in peaking times here is not more than 4 hours. Analysis of data in more conditions using the above approach could inform the search for the molecular players that determine division timing in *S. elongatus*.

**Table 1:** Distinguishing between the modulated rates and the gating models. The models predict different molecular players to implement the effects on division timing by the clock. The table shows the circadian phase at the peak of the bioluminescent reporter under the *kaiBC* promoter measured in related experiments [26], as well as the concentration of the divisor and effector in the modulated rates and gating models, respectively, as determined from the best fit values of *φ* in the two models (SM Section 5.1).

|  | Circadian phase at peak activity (h) |  |  |
| --- | --- | --- | --- |
|  | Related experiments | Modulated rates | Gating |
| LL | - | $12 \pm 1$ | $13 \pm 1$ |
| 16:8 LD | $16 \pm 1$ | $14 \pm 1$ | $11 \pm 1$ |
| 12:12 LD | $14 \pm 1$ | $12 \pm 1$ | $8 \pm 1$ |

**Table 2:**
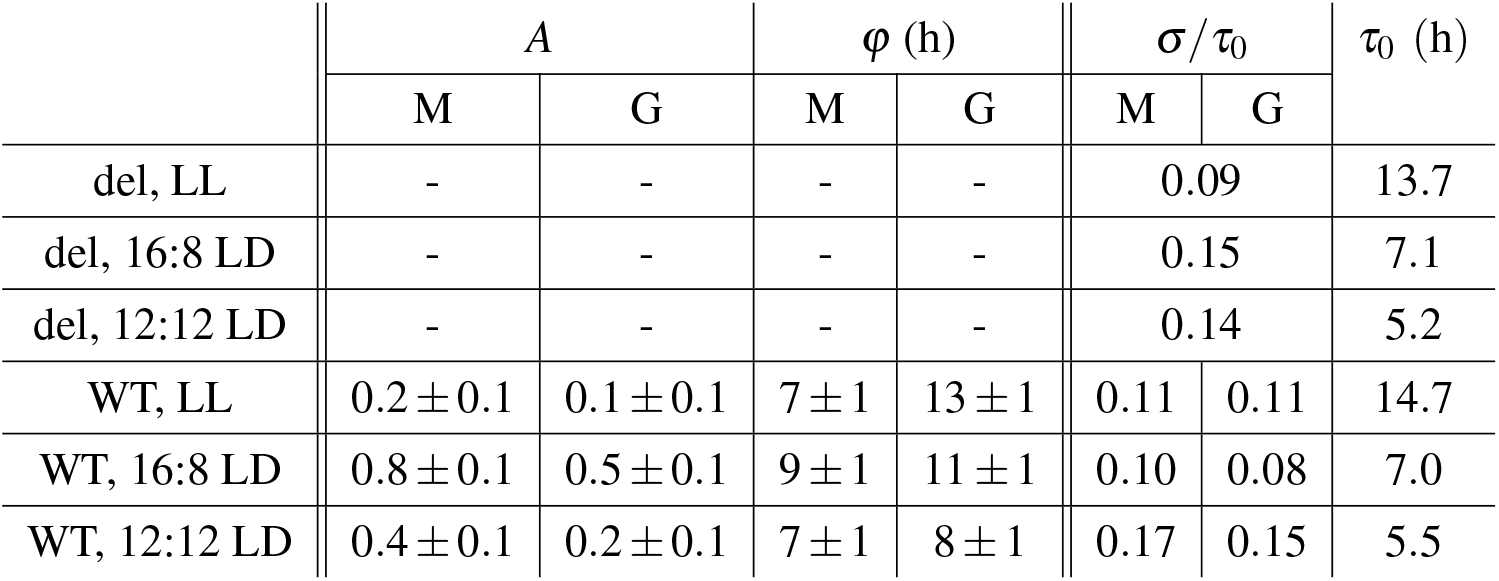
The best fit or extracted values for the parameters of the modulated rates (M) and the gating (G) models for the wild type (WT) and the clock-deletion (del) strains under various environments. - denotes parameters not applicable to that condition. *τ*_0_ = ln 2/*λ*_0_ is the fastest doubling time corresponding to the maximum growth rate *λ*_0_, and is extracted from the experiments. *σ* is chosen to match the experimentally observed standard deviation of cell lengths at birth. The best fit value of *r ≈* 0.025±0.006 h^−1^, corresponding to a half-life of 28±6 h, was determined for the clock-deletion strain under 16:8 LD, and was used for all environments. Error bars on *A*, *φ*, and *r* denote intervals (to the precision specified) outside which the best fit value has a significance level smaller than 0.01 by likelihood analysis (Methods).

**Table 3:** Generation time correlations between mother-daughter, sister-sister, and cousin-cousin pairs (***ρ***_*md*_, ***ρ***_*ss*_, ***ρ***_*cc*_, respectively). The errors on the experimental values report 95% confidence intervals. Table S2 contains the parameter values used.

|  | Wild type under LL | Modulated rates | Model without clock | Clock-deletion under LL |
| --- | --- | --- | --- | --- |
| $\rho_{md}$ | $-0.25 \pm 0.12$ | $-0.46$ | $-0.27$ | $-0.07 \pm 0.13$ |
| $\rho_{ss}$ | $0.62 \pm 0.10$ | $0.68$ | $0.31$ | $0.41 \pm 0.11$ |
| $\rho_{cc}$ | $0.46 \pm 0.10$ | $0.56$ | $0.21$ | $0.29 \pm 0.13$ |

Similarly, the best fit values of *φ* are more parsimoniously interpreted in the modulated rates model. Experiments have shown that for different values of *T_L_* (Eq. 2), the concentrations of the Kai proteins shift in circadian phase such that the phase at the peak increases by *T_L_*/2, or “mid-day tracking” [26]. Consistent with this observation, the best fit value of *φ* under 16:8 LD is two hours more than that under 12:12 LD in the modulated rates model (Table 1). Also in the modulated rates model, the best fit value of *φ* under LL is the same as that under 12:12 LD, consistent with the fact that the clock was entrained under 12:12 LD (Table 1). In contrast, the best fit value of *φ* in the gating model under LL is five hours different from that under 12:12 LD, suggesting that the molecular players in the gating model do not follow the mid-day tracking activity of the Kai proteins. Note, however, that the experiments in Ref. [26] were done with on-off light intensity profiles without the sinusoidal dependence used in Ref. [5]. Therefore, further experiments to determine the activity of the Kai proteins, and other potential modulators of division timing, would help verify the above distinction between the two models.

The differences between the two models in predictions involving *φ* arise from the difference between integrating a signal over time, and acting on the signal instantaneously. By taking the derivative of Eq. 9, the gating model can be rewritten as,

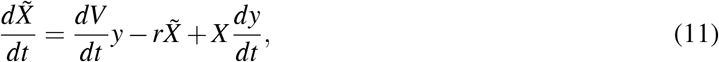

which is equivalent to the modulated rates model for the variable 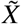 with an extra term *X* (*dy/dt*). When the degradation rate is small compared to the growth rate, as is the case here, *X* approximately scales like *dV/dt*. The extra term therefore approximately modulates the rate of divisor accumulation by both *y* and the derivative of *y*. The form of the extra modulation explains why both models can capture the effects of the clock on division timing, albeit with quantitatively different predictions involving the best fit values of *φ*.

Importantly, the modulated rates model predicts no divisions during darkness, whereas the gating model can lead to divisions during darkness without growth. The latter case occurs when enough divisors have accumulated, but not enough are active according to the gating function *y* (*θ*). Divisions can then occur just by the passage of time, without cell growth, and the consequent activation of divisors with increasing *y* (*θ*). The above scenario can be demonstrated in a numerical simulation using the best fit parameters under 16:8 LD, and tracking the division events for cells entrained under 16:8 LD but imaged during a cycle where the light is turned off abruptly during the day. The gating model predicts that a noticeable fraction of cells will divide during darkness in this scenario, whereas the modulated rates model predicts no divisions during darkness (Fig. 4a). Divisions in darkness have indeed not been observed experimentally. However, it may be that cells possess additional mechanisms to abort divisions during darkness, regardless of how the clock affects division timing. One way to distinguish between the two models while circumventing this possibility is to decrease the light intensity abruptly to a small but non-zero value. In this case, the gating model predicts that a larger fraction of cells will divide afterwards (Fig. 4b). We note the caveat that the clock will likely be re-entrained by the abrupt change in light intensity, and hence, *y* (*θ*) will be affected on longer time scales. Nevertheless, on the shorter time scale shortly after the change in light intensity, our predictions will hold. The above difference between the two models could be relevant for cells in nature facing fluctuations in environmental light intensity [29]. The experimental realization of the scenario would be one way to directly differentiate the two models.

**Figure 4:**
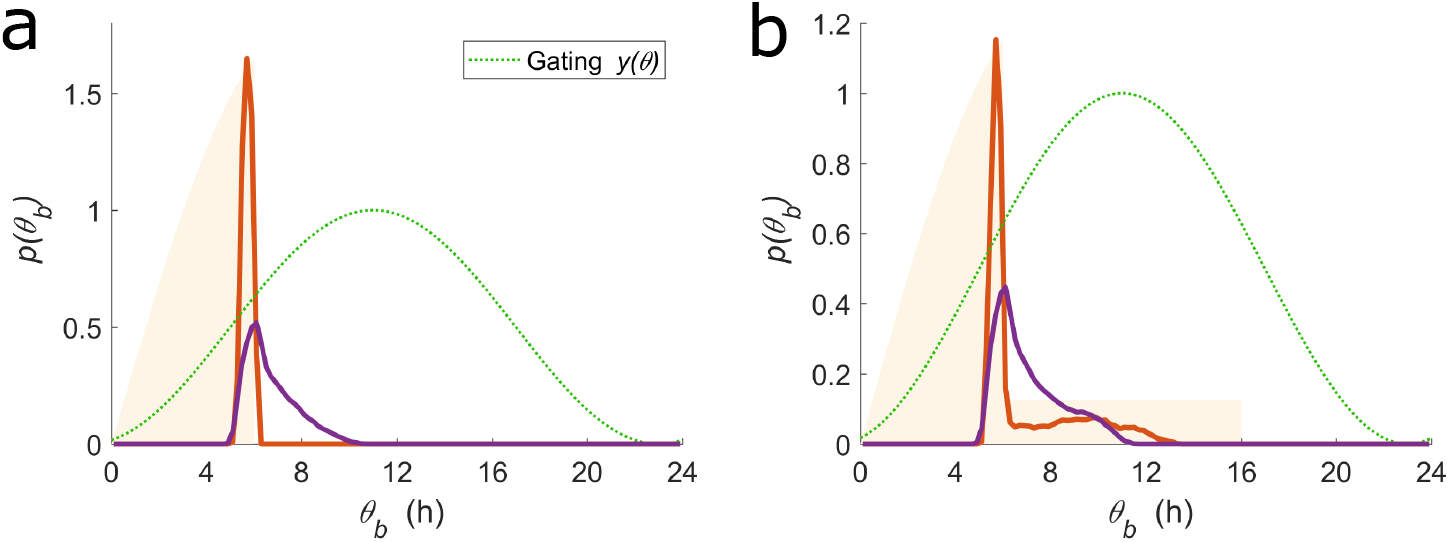
The modulated rates model robustly places divisions away from darkness, whereas the gating model does not. Predictions of the modulated rates (red) and gating (purple) models entrained under 16:8 LD and imaged for one cycle where the light is abruptly turned off (a) or down (b). The simulations used *y* (*θ*) best fit to the data of Ref. [5]. The *y* (*θ*) in the gating model is shown in the green dotted line. Yellow shade shows the light profile during the imaging cycle.

## 3 Discussion

How cyanobacteria regulate division timing has been studied for decades, but how and why the clock regulates division timing remain unclear [2, 4, 5, 30]. One widely considered mechanism is that of gating, where the signal from the clock is assumed to suppress divisions in a specific time window [2, 3]. Here, we proposed a different mechanism of modulated rates, where the signal from the clock is integrated over time to affect division timing. Biologically, the gating model could correspond to a post-translational mechanism while the modulated rates model could correspond to a transcriptional mechanism.

To distinguish between the two mechanisms, we formulated a simple framework that describes how cell volume growth, the environmental light profile, and the internal circadian clock together determine division timing. Our framework differs from existing ones in both formalism and structure. Ref. [4] modeled the relation between the progression of the circadian phase and that of division timing with a general nonlinear map. Ref. [30] studied a model in which the generation time is determined by a linear combination of the previous generation time and an oscillatory function of the circadian phase. The above approaches did not consider the feedback of cell size on division timing. However, for exponentially growing cells such as those of *S. elongatus*, timing divisions without feedback from cell size fails to maintain a homeostatic average cell size [7]. Ref. [5] accounted for the effects of cell size regulation by modeling the instantaneous probability to divide as a function of cell size multiplied by the growth rate and a periodic coupling function of the circadian phase [31]. The approach of Ref. [5] can describe the experimentally observed statistics of division timing under LL. However, the coupling function fitted to LL data cannot capture the low density of divisions in the early hours of the light period under LD (SM Section 5.5). It is also not straightforward to parametrize the coupling function to gain an understanding of the underlying molecular mechanism, which will require further work. Our models specify division timing via simple deterministic dynamics and implement stochasticity via a coarse-grained noise term [14]. The simplicity provides mechanistic insights by describing how the clock affects division timing via two parameters with mechanistic interpretations.

With our framework, the modulated rates model appears to be more consistent with existing experiments than the gating model. Moreover, existing data is consistent with the simple mechanism that division timing is regulated by the accumulation of a division limiting protein in phase with genes whose activity peak at dusk. Suggestively, FtsZ is one such protein [25]. Together with further single-cell level experiments, our simple and illustrative modeling framework will be useful in unraveling how the clock regulates division timing.

## 4 Methods

### Numerical simulations of the models

The deterministic generation time was determined by numerically integrating the equations for the accumulation of divisors, Eqs. 3 or 7. The stochastic generation time is obtained via Eq. 6. Cell volume is calculated according to Eqs. 1–2 and is divided in half at division. The process is repeated for at least 10^5^ generations, following only one of the newborn cells at division. To describe the distribution of circadian phases at birth under LL, a similar method was used to track division events of a growing colony (SM Section 5.3).

### Determination of the best fit values for model parameters

The best fit value of *r* in Eq. 3 was determined as follows. For a given *r*, *σ* in Eq. 6 was chosen to match the CV of *l_b_*. Then, the best fit value of *r* for the clock-deletion strain under 16:8 LD was chosen to minimize the sum of squared residues between model predictions and experimental observations for *p*(*θ_b_*) and *p*(*t_d_*), with bin size corresponding to the experimental time resolution (0.75 h under LL and 1 h under LD). The best fit values of *A* and *φ* for the wild type strain under LD were determined by minimizing the same quantity. For the wild type strain under LL, the best fit values were chosen to minimize the sum of squared residues in the correlations between *t_d_* and *θ_b_*, binned according to *θ_b_* with bin size corresponding to the experimental time resolution. Table S2 summarizes the best fit values of all parameters obtained.

### Determination of the goodness of fit

The goodness of fit of the models and the errors on the best fit values of the model parameters can be estimated by comparing the residue between the best fit predictions (best residue) and that between the predictions of the divisor accumulation model without a clock (worst residue). To determine the error bars on the best fit values, we held other parameters constant and scanned the parameter in question until the residue becomes larger than 5% the difference between the best and worst residue. We determined error estimates to the decimal place for *A*, and to the hour for *φ* and the half-life corresponding to *r*.

## Acknowledgments

PH was supported by the Quantitative Biology Initiative Student Award, the Aramont Fund for Emerging Science Research, and the NSF MRSEC DMR-1420570. BMCM was supported by the UK Biotechnological and Biological Sciences Research Council Synthetic Biology Research Centre “OpenPlant” Award BB/L014130/1. AA thanks support from NSF CAREER 1752024 and the Harvard Dean’s Competitive Fund.

## 5 Supplementary materials

### 5.1 Properties of the model

In this section, we provide details of various aspects of the modulated rates model that pertained to our analysis of the model against experiments. We first elaborate on our motivation to investigate the divisor accumulation model. We then provide intuition for various behaviors of the model. Finally, we calculate the divisor concentration within the modulated rates model.

#### Motivation for the divisor accumulation model

Without degradation, i.e. *r* = 0 in Eq. 3, the divisor accumulation model reduces to an “adder” model, where cells on average add a constant size increment from birth to division [19]. This fact and the observation that the clock-deletion strain under LL behaved approximately like an adder motivated us to investigate a divisor accumulation model. However, we found that the divisor accumulation model without degradation cannot capture the full extent of the bias of divisions away from dawn (Fig. S1). Moreover, it also cannot capture the mean generation times of the short- and long-generation cells (Fig. S1). Instead, we found that a degradation rate *r* ≈ ln2/(28 ± 6 h) ≈ 0.025 ± 0.006 h^−1^ is required to capture these statistics of division timing for the clock-deletion strain under 16:8 LD (Fig. 2c and Fig. S1).

**Figure S1:**
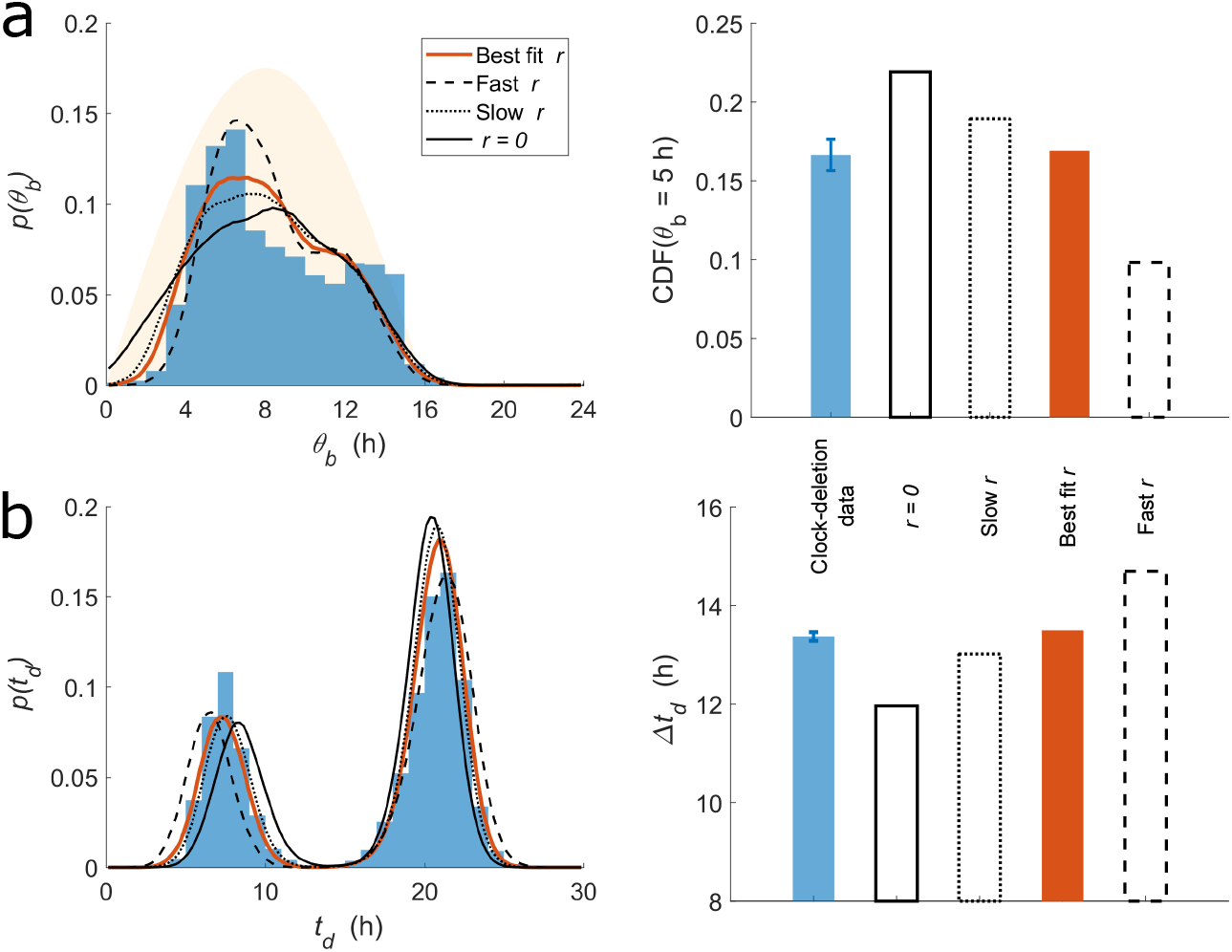
The effects of degradation rate on the distributions of *θ_b_* and *t_d_*. Blue denotes data of the clock-deletion strain under 16:8 LD. Red denotes the divisor accumulation model with the best fit value of *r*. The black lines show model predictions for different values of *r*, corresponding to half-lives of 44 (dotted) and 12 (dashed) hours. The solid black line shows model predictions for *r* = 0. The bar plots show the cumulative fraction of divisions that have occured before the specified circadian phase CDF(*θ_b_* = 5 h), and the difference between the mean generation times of short- and long-generation cells, Δ*t_d_*.

#### Intuition for model behavior

Compared to the case without degradation, the long-generation cells now have even longer generation times due to the degradation of accumulated divisors during darkness. The long-generation cells must now compensate for the degraded divisors, and therefore will be larger at division. Cells with the largest *t_d_* are therefore also the largest cells at division (Fig. S2a). These cells might go on to become short-generation cells with small generation times. In this way, degradation increases the difference between the mean generation times of short- and long-generation cells. Similarly, coupling division timing to the clock introduces correlations between cell size and division timing, as explained in the main text and shown in Fig. S2b, as well as increases the difference between the mean generation times of short- and long-generation cells.

**Figure S2:**
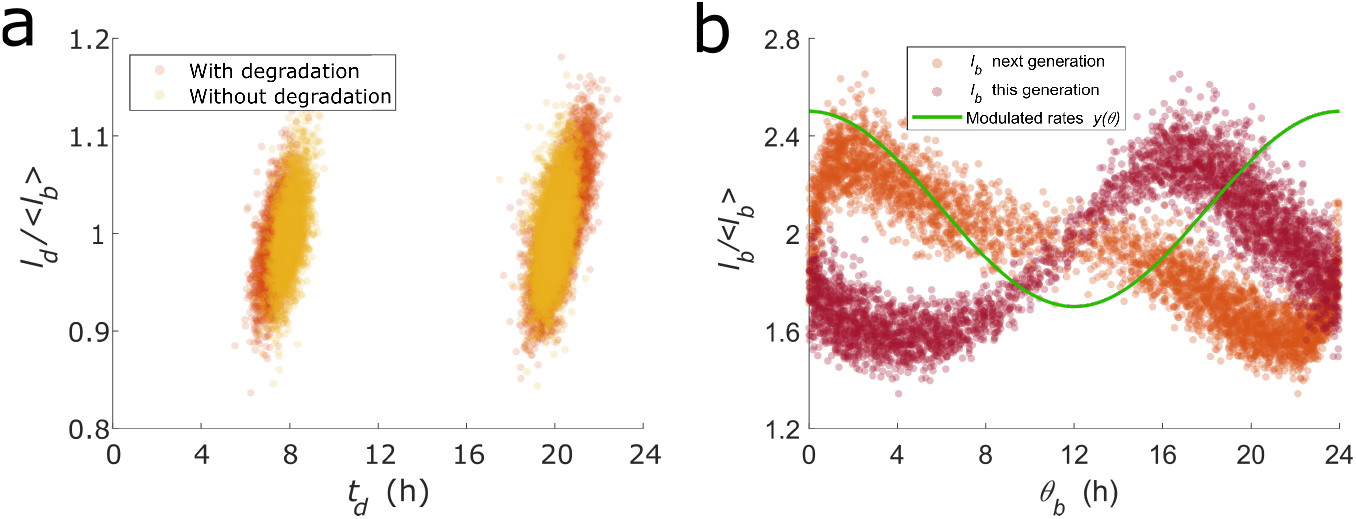
Predictions of the modulated rates model with parameters chosen to highlight the origins of the observed correlations between cell size and division timing. (a) The basic model with *τ* = log(2)/*λ* = 8 h, under on-off 12:12 LD, and with *r* = log(2)/20 h^−1^ (red) and without degradation (yellow). Cells with large *t_d_* are also larger in size at division. (b) The modulated rates model with *τ* = 10 h, under constant light, and *A* = 0.4. For example, a large cell born approximately 16 h into the day will likely become a smaller than average cell in the next generation, leading to correlations between cell size and division timing.

#### Divisor concentration

Within the modulated rates model, the concentration of the divisor, *x* = *X/V*, lags some time behind the promoter activity described by *y* (*θ*). The lag duration can be estimated by ignoring divisions and calculating the resulting concentration by integrating for *X*(*t*)/*V*(*t*) as *t* → ∞, which gives

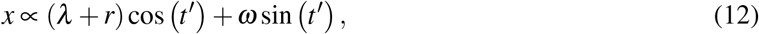

where *t*′ = *ωt* + *πφ*/12. Therefore, if (*λ* + *r*) ≫ *ω*, then *x* ∝ *y* and the promoter activity and the concentration are approximately synchronous. If (*λ* + *r*) ≪ *ω*, then the concentration lags a quarter of a period behind. The values of *λ* and *r* for the experiments we analyzed lie approximately in the latter regime. More precisely, the concentration lags approximately 5±1 hours behind the promoter activity. Together with the best fit value of *φ* in Table S2, the circadian phase at the peak of the divisor concentration in Fig. 4c can then be obtained.

### 5.2 Summary of the parameter values used and comparisons to data

For the purpose of analyzing the statistics of division timing, there are only two free parameters (*A* and *φ*) in the models, whereas the other parameters (*λ*_0_, *r*, *σ*) are already determined and fixed or extracted from the data. The maximum growth rate *λ*_0_ is extracted from the experimentally observed instantaneous growth rates. As explained in the main text, the degradation rate *r* is assumed to be the same as inferred from the experiments with the clock-deletion strain. The magnitude of the coarse-grained stochasticity *σ* is chosen to match the experimentally observed CV of cell lengths at birth. The two remaining parameters describing the modulation function *y* (*θ*) must then explain the statistics of division timing, including those shown in Figs. 2, 3, Fig. S3, and Table S3.

**Figure S3:**
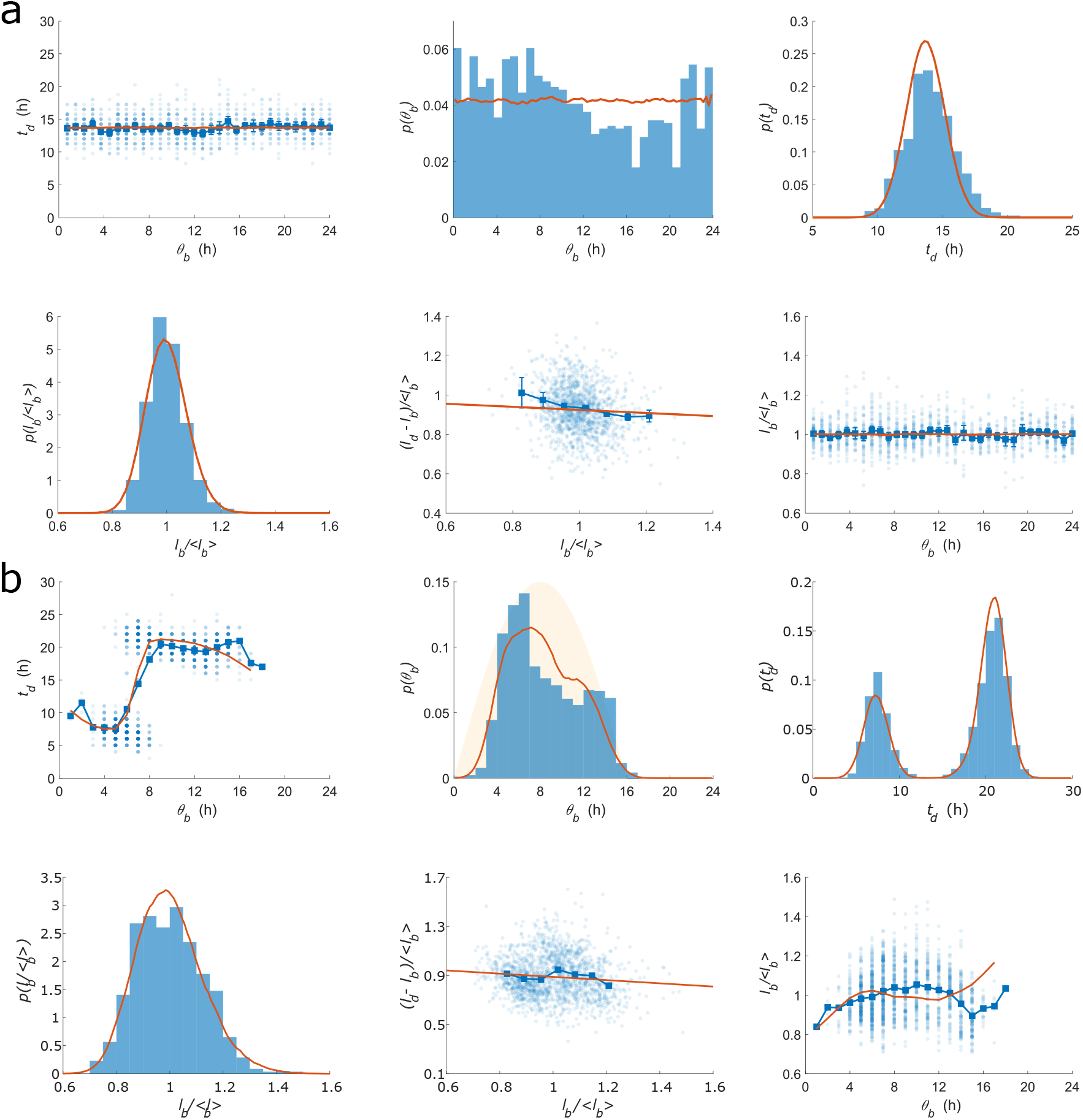
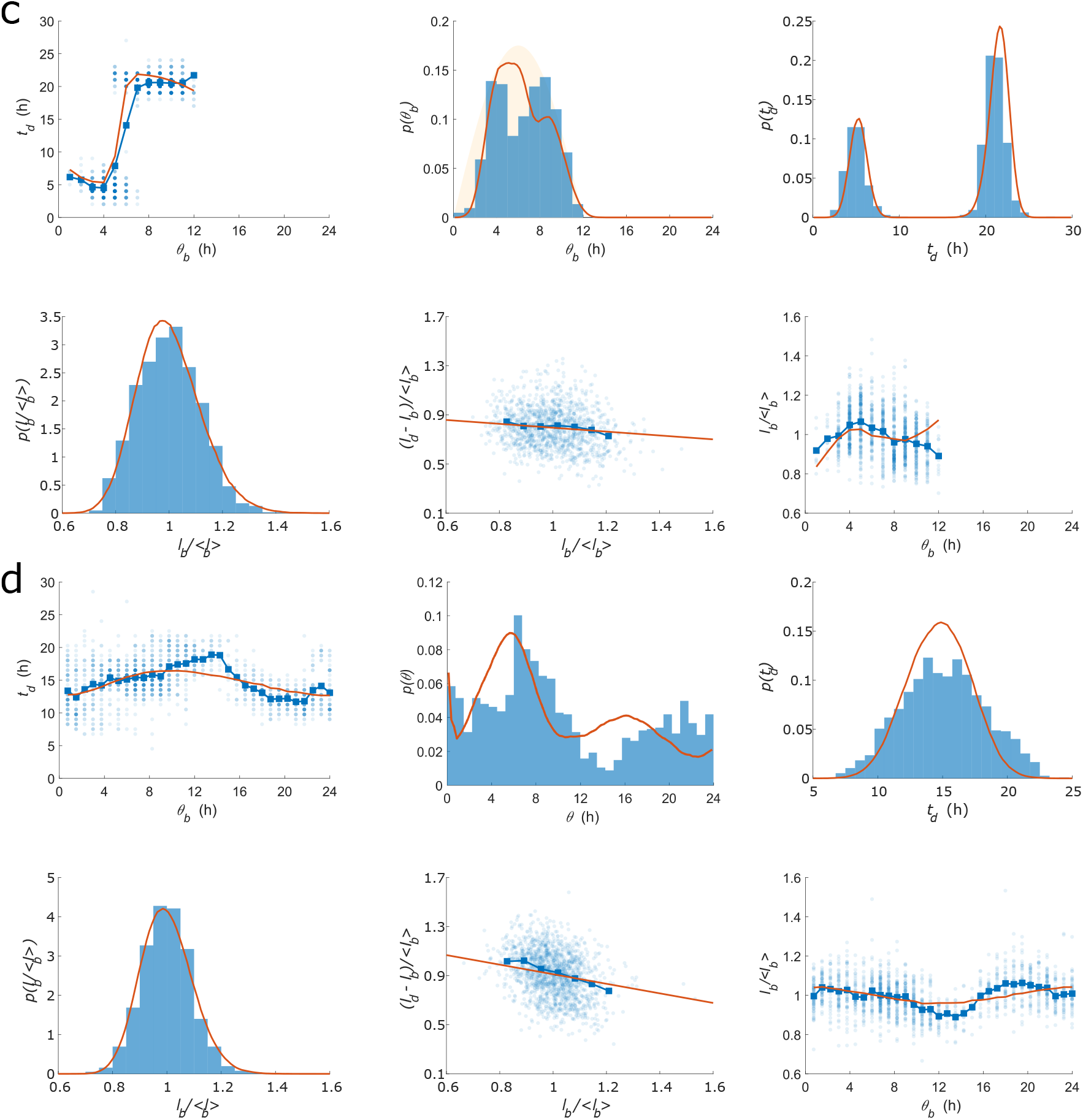
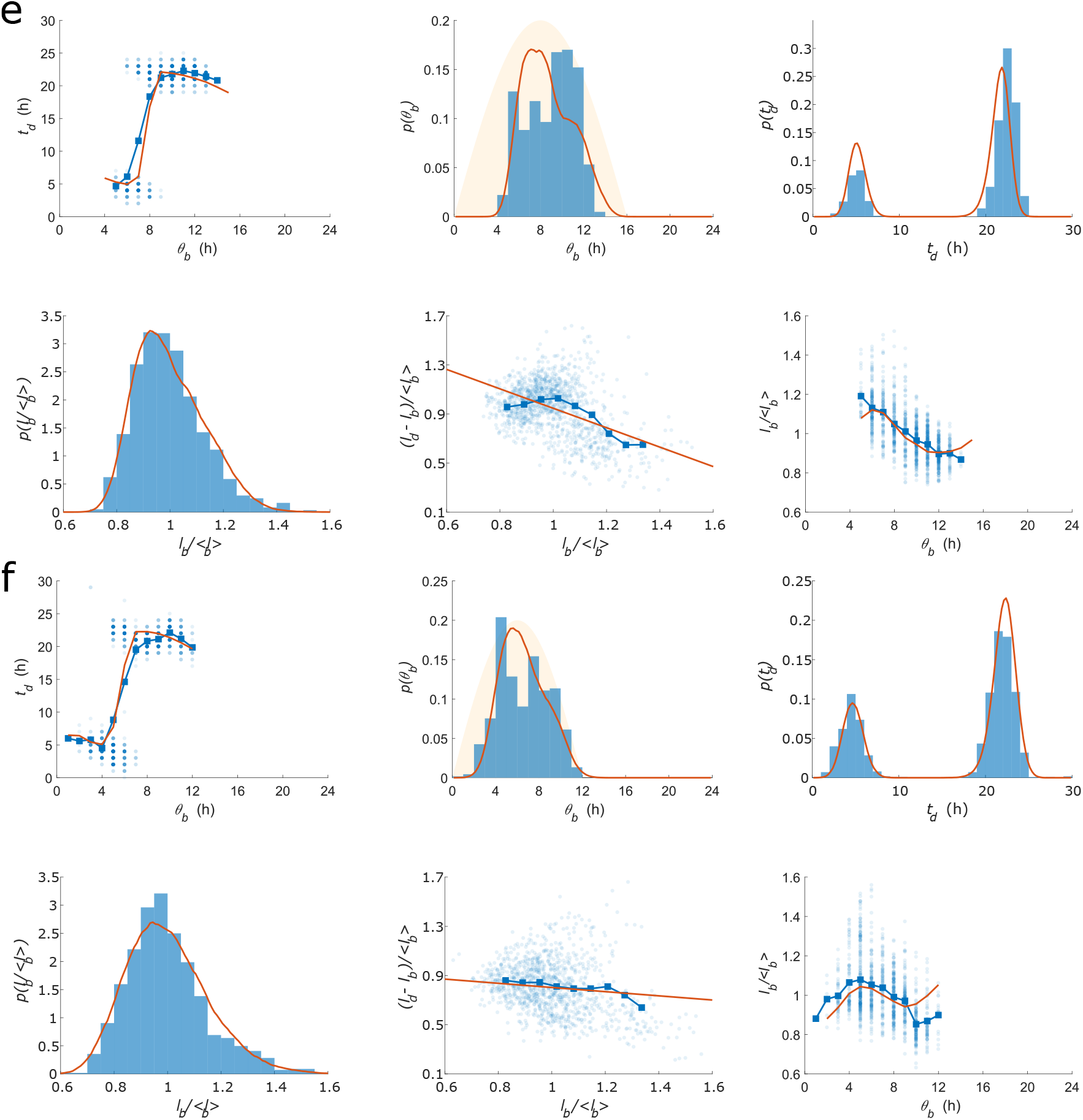
Divisor accumulation with modulated rates captures the statistics of division timing in *S. elongatus*. Panels show model predictions compared with experiments of the clock-deletion strain (a-c) and the wild type strain (d-f) under LL (a,d), 16:8 LD (b,e), and 12:12 LD (c,f). Figure legends are the same as Fig. 3 The variables are defined in the caption of Fig. 1. Table S2 contains the parameter values used.

**Figure S4:**
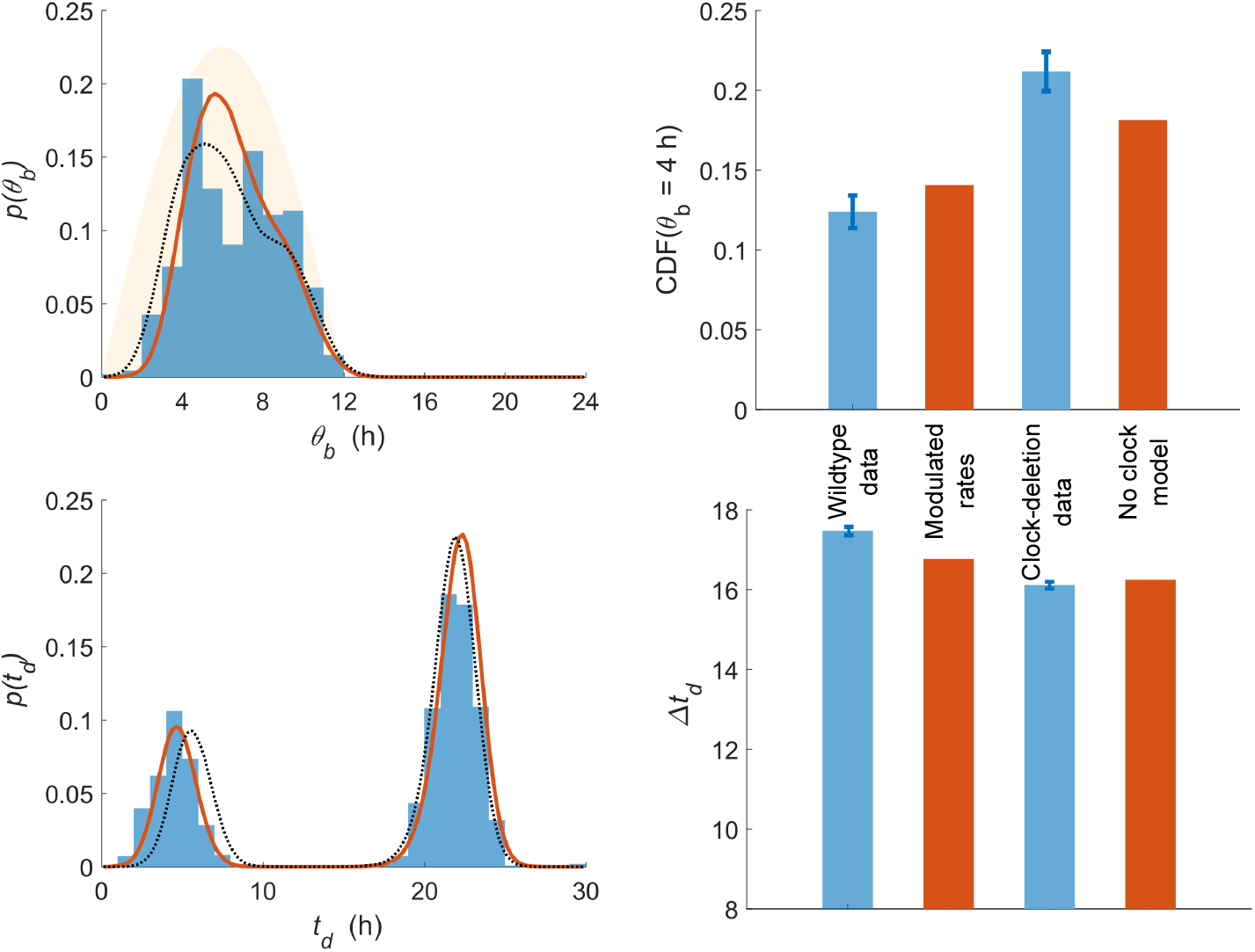
Divisor accumulation with modulated rates can describe division timing in the wild type strain under 12:12 LD. Figure legends and axes labels are the same as in Fig. 3.

**Figure S5:**
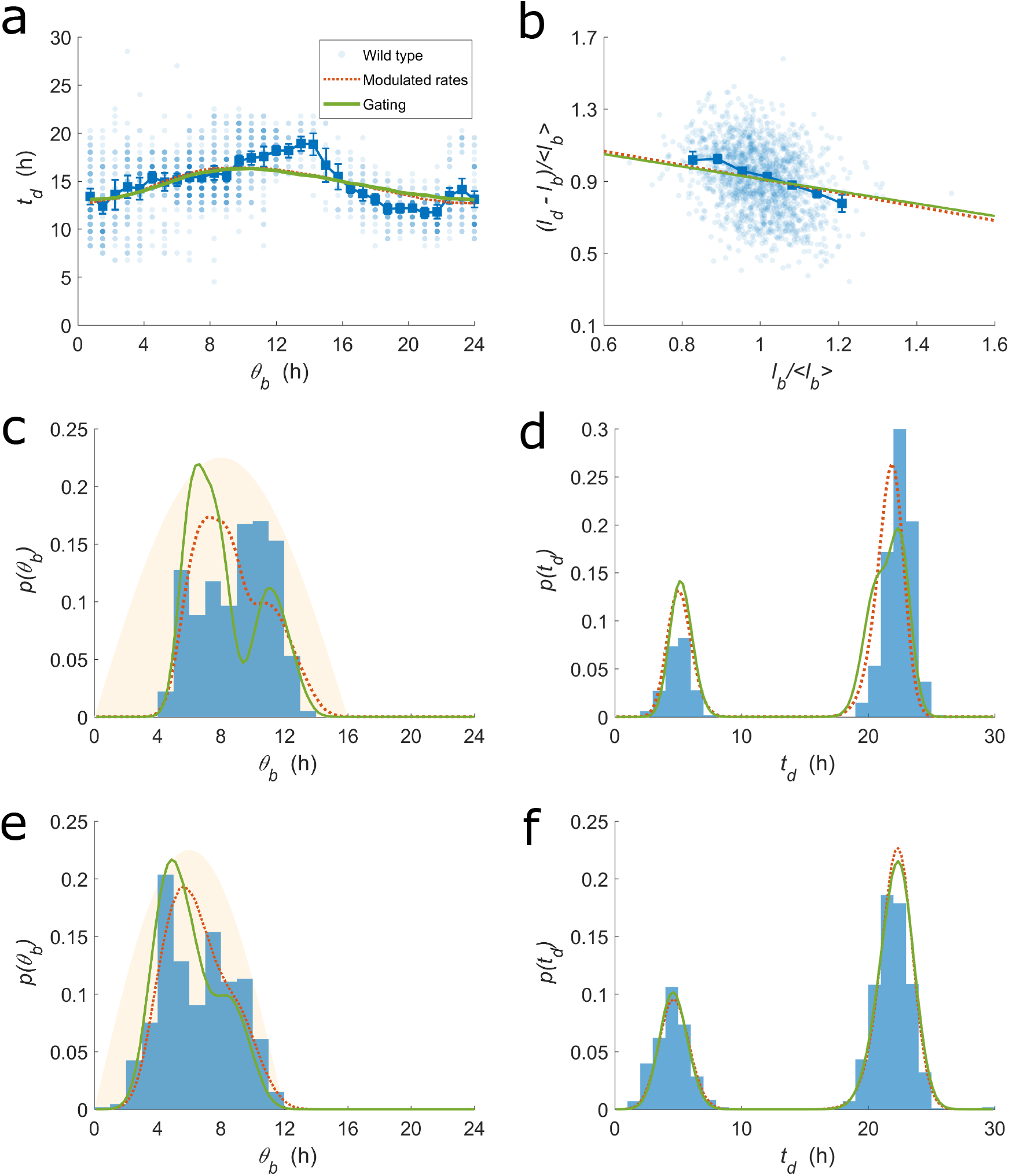
Divisor accumulation with gating can describe division timing in the wild type strain. Figure legends and axes labels are the same as in Fig. 3, except that green lines here denote predictions of the gating model.

### 5.3 The effects of the statistical ensemble

The details of the experimentally recorded ensemble affects the reported statistics of division timing. The recorded cells may be sampled from the entire population tree or a few lineages, from experiments that begin with synchronized or asynchronized cells, or from experiments that end at different times during the day. Such details significantly affect the shape of *p*(*θ_b_*) under constant light conditions. For statistics from a single lineage, the two distributions *p*(*θ_b_*) and *p*(*θ_d_*), where *θ_d_* = (*θ_b_* + *t_d_*) mod 24, are exactly the same, since the circadian phase at division is simply the circadian phase at birth for the next generation in the lineage. However, for synchronized cells in a growing population, the two distributions could be drastically different depending on when the experiments end, since the cells are synchronized and will tend to divide nearby in time. The modulated rates model, when simulated at the population level while taking into account the recording protocol (i.e. recording divisions of growing populations of synchronized cells, beginning at dawn, for 96 hours), reproduces *p*(*θ_b_*) and *p*(*θ_d_*) in qualitative agreement with experimental observations (Fig. S6a). The above effect does not significantly affect the features of *p*(*θ_b_*) under LD. When divisions begin to occur, as quantified by CDF(*θ_b_* = 5 h), are more similar between single-cell (i.e. Methods, Numerical simulations of models) and population level simulations than between simulations with different values of *φ* (holding all other parameters constant), implying that our fitting procedure can determine the best fit value of *φ* to the hour (Fig. S6b).

**Figure S6:**
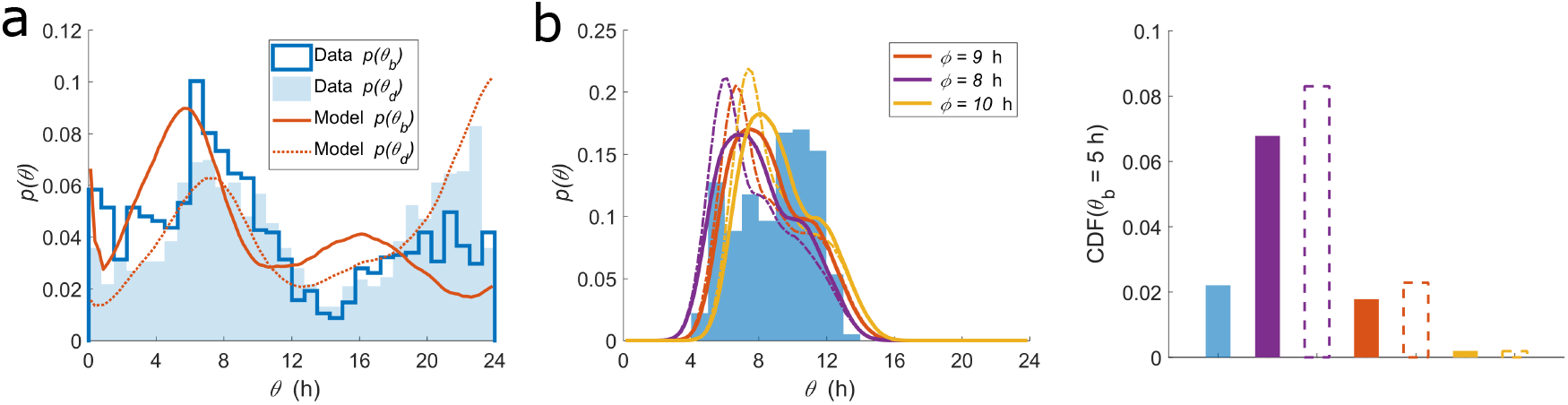
The distributions of circadian phases at birth and at division can be captured, after taking into account the details of the statistical ensemble. (a) The experimentally observed (blue histograms) and predicted (red lines) distributions of circadian phase at birth (dark/solid) and division (light/dashed) under LL. The predictions were obtained by simulating the modulated rates model using the best fit values in Table S2 and taking into account the recording protocol. (b) The features of *p*(*θ_b_*) under LD are not significantly affected by the details of the statistical ensemble. Blue histogram shows data for wild type under 16:8 LD. The solid and dashed colored lines shows the predictions of the modulated rates model using single-cell and population level simulations, respectively, for different values of *φ* (holding other parameters constant).

### 5.4 Cousin-cousin correlations

Correlations in generation times along cell lineages have been used to study systems from cyanobacteria to cancer cells [22, 30, 32]. In particular, the relations between the parent-child, sibling-sibling, and cousin-cousin correlations in generation times (***ρ***_*md*_, ***ρ***_*ss*_, ***ρ***_*cc*_, respectively) inform how division timing might be inherited from the parent cell or affected by an underlying clock-like process. The modulated rates model captures these correlations (Table S3).

### 5.5 Further comparisons of the modulated rates model

To better understand the modulated rates model, we compared it to the model of Ref. [5]. First, we calculated the likelihood functions for the model of Ref. [5]. Briefly, the model assumed that the instantaneous probability to divide is dependent on the instantaneous cell size, cell size at birth, the growth rate, and the circadian phase. In particular, the circadian phase modulates the probability to divide by a coupling function (*G*(*t*) in Eq. 1 of Ref. [5]) that is independent of the environment. We found that under LL conditions, the modulated rates model and the model of Ref. [5] have comparable goodness of fit to the data (Fig. S7a). However, under LD conditions, the model of Ref. [5] does not capture the full extent of the bias of divisions away from darkness under 16:8 LD (Fig. S7b). This result suggests that within the framework of Ref. [5], the coupling function either varies with the environment or is modulated by an extra environmental factor, analogous to our conclusion that within the modulated rates model, the best fit values of *A* and *φ* vary with the environment.

**Figure S7:**
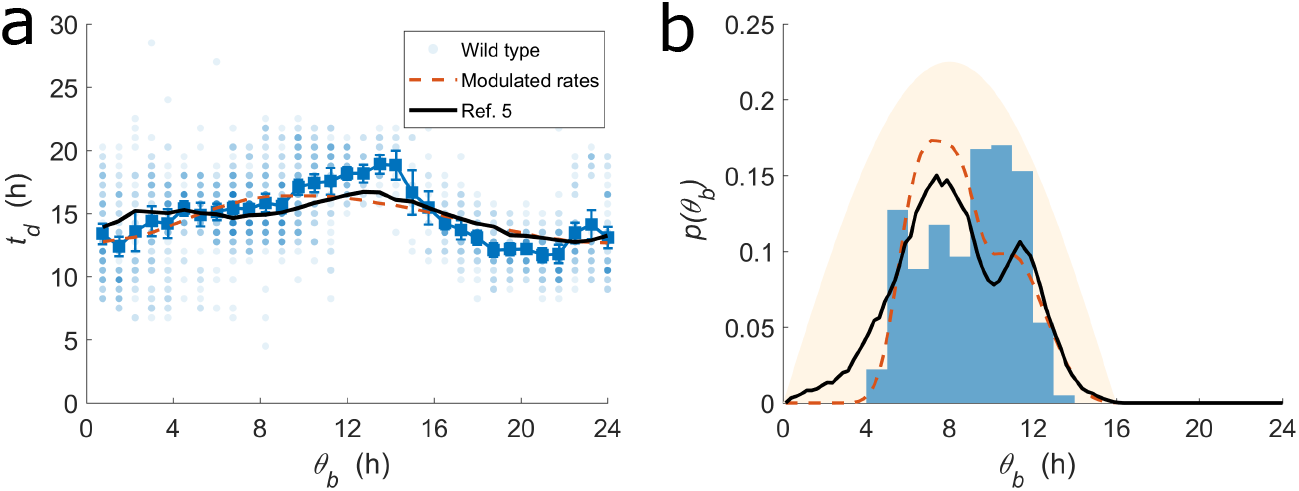
Comparisons of the modulated rates model to the model of Ref. [5].

